# High standing genetic variation in an invasive plant allows immediate evolutionary response to climate warming

**DOI:** 10.1101/2020.02.20.957209

**Authors:** Yan Sun, Oliver Bossdorf, Ramon Diaz Grados, ZhiYong Liao, Heinz Müller-Schärer

**Author notes:** Department of Biology/Ecology & Evolution, University of Fribourg, Chemin du Musée 10, CH-1700 Fribourg, Switzerland. The name and complete mailing address (including telephone and fax numbers and e-mail address) of the person to whom correspondence should be sent: Yan Sun, University of Fribourg, Dept. of Biology, Chemin du Musée 10, CH-1700 Fribourg, Switzerland. Emails of all authors Yan Sun, Oliver Bossdorf, Ramon Diaz Grados, ZhiYong Liao, Heinz Müller-Schärer. Authorship Y.S., H.M.S. and O.B. designed the experiment, Y.S. and H.M.S. conducted the field experimental evolutionary study, Y.S., R.D.G. and Z.Y.L. conducted the common garden experiment, Y.S. performed all genomic and statistical analyses, Y.S. wrote the first draft of the manuscript, Y.S., H.M.S. and O.B. contributed substantially to the final revision.

## Abstract

Predicting plant distributions under climate change is constrained by our limited understanding of potential rapid adaptive evolution. In an experimental evolution study with the invasive common ragweed, we subjected replicated populations of the same initial genetic composition to simulated climate warming. Pooled DNA sequencing of parental and offspring populations showed that warming populations experienced a greater loss of genetic diversity, and greater genetic divergence from their parents, than control populations. In a common environment, offspring from warming populations showed more convergent phenotypes in seven out of nine plant traits, with later flowering and larger biomass, than plants from control populations. For both traits, we also found a significant higher ratio of phenotypic to genetic differentiation across generations for warming than for control populations, indicating stronger selection under warming conditions. Our findings demonstrate that ragweed populations can rapidly evolve in response to climate change within a single generation.

## Introduction

There is overwhelming evidence that the global climate is changing rapidly^1^. Besides a rapid increase in global temperatures, the frequency of extreme weather events is also increasing, often confronting species with environmental conditions they rarely experienced in their recent evolutionary histories^2^. Most previous research has focused on range expansion or contraction due to changes in environmental settings, while evolutionary responses to climate change are still understudied, even though it is well known that historical climate changes have often been accompanied by evolutionary changes^3,4^. In parallel to climate change, biological invasions, another important driver of ecological changes in the Anthropocene^5^, are also increasingly threatening native ecosystems, with no saturation yet in the accumulation of alien plants worldwide^6^ and thus with increasing costs for their management^7^. Theoretical and empirical studies indicate that climate change may exacerbate the risk of plant invasions^8^, but understanding the interactive effects of climate change and invasions remains challenging, particularly since invasive populations are often also evolving rapidly.

Biological invasions are often accompanied by significant demographic or evolutionary events such as genetic bottlenecks, hybridization and admixtures, and therefore invasive populations are often not representative of the genetic diversity in their native ranges^9^. Moreover, there is often intraspecific hybridization in invasive populations. Population admixtures before introductions or after multiple introductions from genetically distinct sources have repeatedly been found to overcome initial demographic bottlenecks, thereby solving the so-called “genetic paradox of invasion^10^, the invasion success of some species despite strong demographic bottlenecks. In a recent review, Dlugosch et al.^10^ reported admixture of 37% of 70 invasive species studied with nuclear markers. Such admixture can increase genetic variance within populations, and it can result in heterosis, as well as novel and/or transgressive phenotypes^11,12^. Moreover, escape from specialized natural enemies and mutualists may result in a simpler suite of interactions for invasive plants in their introduced range^13^, which could relax selection on multiple traits, allowing populations to adapt more rapidly to novel environmental conditions than native species.

Microevolutionary change in plants can indeed be fast^14^, with several examples in particular from climate change experiments and invasive plant populations^15^. Recent studies found, for instance, rapid molecular divergence in plants subjected to experimental climate change^16^, usually with reduced genetic variation under selection (cf. review Hoffmann and Sgrò^17^). Other studies showed rapid life-history differentiation of *Lythrum salicaria* populations in response to latitudinal variation^18^ and warming selection^19^, and rapid differentiation of root:shoot ratio and water-use efficiency among invasive *Solidago canadensis* populations^20^.

In order to predict future ecological consequences of plant invasions, a better understanding of the pace and extent of their evolutionary responses to climate change is crucial^21^. As suggested above, evolutionary responses to climatic change might be even stronger in introduced than in native populations, being further fostered by their relative isolation from gene flow from the native range, as compared to spreading native population, and we may thus expect evolutionary changes influencing range expansion, niche differentiation and ecological impact in invasive plant populations^22,23^.

Microevolution of plant populations in response to environmental change has so far mostly been studied in two ways: First, researchers have measured heritability of target traits as well as contemporary selection on them, and then made predictions about the potential for short-term evolution of traits^24^. Second, other studies compared populations or species that have diverged over time and made inferences about the processes that drove evolutionary change in the past, such as in biological invasion^25,26,27^. A third, and so far less used, approach often considered the “gold standard” of microevolutionary studies are selection experiments – also called experimental evolution - where microevolution is observed “in action”but see Agrawal et al.^28^. Experimental evolution studies can follow evolutionary dynamics under different selection regimes, and assess the repeatability (and thus predictability) of evolution across replicated experimental populations^28,29^. To improve a mechanistic understanding of evolutionary changes, de Villemereuil et al.^30^ proposed combining population genomics with a common-garden approach. Using this approach, Barghi et al.^31^ recently combined a laboratory selection experiment to detect selection signatures with phenotyping in a common environment and showed that *Drosophila simulans* populations harbour a vast evolutionary potential for future temperature adaptation. So far, experimental evolution studies have rarely been done in natural settings^29,32^.

One of the key traits associated with plant responses to environmental stress is specific leaf area (SLA), which is determined by a trade-off between cell size and cell density^33^. For instance, Hudson et al.^34^ found increased SLA in a long-term artificial warming experiment with *Cassiope tetragona*. Another important trait in the context of climate change responses is relative water content (RWC), an indicator of plant water deficit and of the capacity of a plant to avoid dehydration under different growing conditions^35^. Under water deficit conditions, there is often a negative relationship between SLA and RWC, with lower SLA reflecting thicker leaves that can maintain a higher RWC^36^. Net assimilation rate (NAR), a proxy for a plant’s efficiency in using CO_2_ for dry matter accumulation, was found to significantly differ in *Saccharum officinarum* between control and heat stress over time and space^37^.

Here, we studied one of the most noxious plant invaders in Europe, common ragweed *Ambrosia artemisiifolia* L. (Asteraceae) (hereafter: ragweed), causing some 13.5 million people to suffer from ragweed-induced allergies in Europe, causing economic costs of approximately Euro 7.4 billion annually^38^. Species distribution models predict a northward spread of ragweed under climatic change both for the introduced European, East Asian^39,40^ as well as its native North American range^41^. In general, populations may either migrate to follow suitable environmental conditions in space without evolving, or they might locally adapt to novel climatic conditions, with or without migrating^42^. However, even during migration, plants may experience new abiotic and biotic conditions that could exert selection and thus also migrating populations are expected to evolve. Although all of these processes will affect plant distributions under climate change, evolutionary changes have so far been largely ignored in species distribution models^4^. Ragweed is a wind-pollinated and obligate out-crossing annual, with outcrossing rates of 0.93-1.0 recorded in both native and introduced populations^27,43^. Moreover, previous studies found high genetic variation within introduced ragweed populations in Europe^44,45^, most likely because of multiple introduction and pre-admixtures^46^. This resulted in very large effective population sizes, substantial standing genetic variation in ecologically important traits, and novel genetic substrate, as compared to native populations, and on which selection can act and potentially generate lineages with greater or shifted ecological amplitudes or fitness^11,47^.

To test the potential for rapid evolutionary response to climate change, we studied the first offspring generation from an ongoing experimental evolution study established in 2016 in which replicated populations of identical initial genetic composition had been subjected to simulated climate warming or ambient (control) climate conditions. We employed population genomic tools (pooled DNA sequencing) to test for genomic changes, and a common-environment study using seeds from all parental and offspring populations to assess evolutionary changes in phenotypes. Combining comparisons both between generations (allochronic) and between treatments (synchronic) (cf. Fig. 1, Material & Methods) allowed to specifically ask (1) whether there were significant molecular changes across generations, and if these differed between warming and control populations, (2) whether such molecular changes were accompanied by changes in phenotypic traits, and (3) if so, whether a comparison between phenotypic and molecular changes indicated phenotypic selection and thus rapid genetic response to selection.

**Fig. 1.**
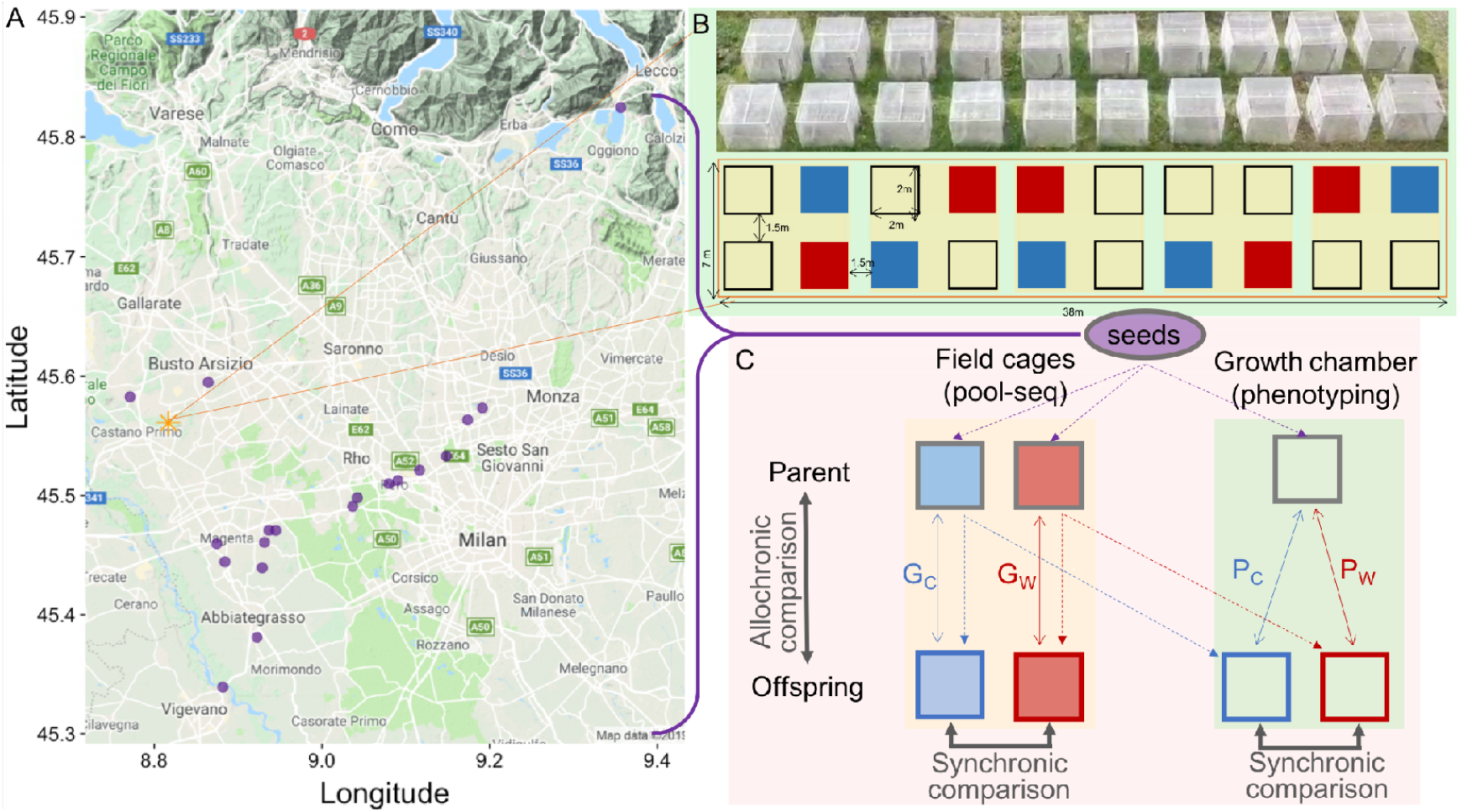
(A) The 19 populations of origin (purple points, in radius of 35 km) of the 60 maternal families of *Ambrosia artemisiifolia* used in our field experiment. The orange star represents the field experiment location. (B) The experimental set-up, with blue (control treatment) and red (warming treatment) cages used in this study (the black frames indicate another treatment not part of this study), each yellow block contains one pair of the two treatments. (C) Overview of the experimental generations and possible comparisons (allochronic and synchronic), with grey frames for populations established from the parental seeds, and blue and red frames for the populations established by the offspring seeds from control and warming treatments, respectively. The blue, red and transparent color filled in the block represent the field control and warming treatments assessed in the common conditions of growth chambers. Dash arrows indicate use of seed sources, and solid arrows are comparisons of molecular or phenotypic comparisons made between generations, with the G_C_ and G_W_ referring to the rates of sequence divergence between generations, and P_C_ and P_W_ referring to the rates of phenotypic divergence between generations (see details in Materials and Methods).

## Results

Based on pool-seq data, the PCA visualising the genetic similarities of all parent and offspring populations showed a much stronger separation between the two generations under warming conditions than under control conditions, and with more similar among populations within both offspring control and offspring warming ellipses as compared to their respective parental populations (Fig. 2A). Moreover, in the warming treatment offspring populations had a significantly lower Tajima’s π (genetic variability within population) than their parental populations (*t* = -3.02, df = 32, *P* = 0.005; Fig. 2B). In contrast, there were no significant differences in Tajima’s π between the two generations in the control treatment (*t* = 1.29, df = 32, *P* = 0.21; Fig. 2B). The cross-generation *F*_ST_ values were significantly higher for warming populations than for control populations (χ^2^ = 4.29, *P* = 0.04; Fig. 2C), indicating stronger genetic divergence under warming conditions.s

**Fig. 2.**
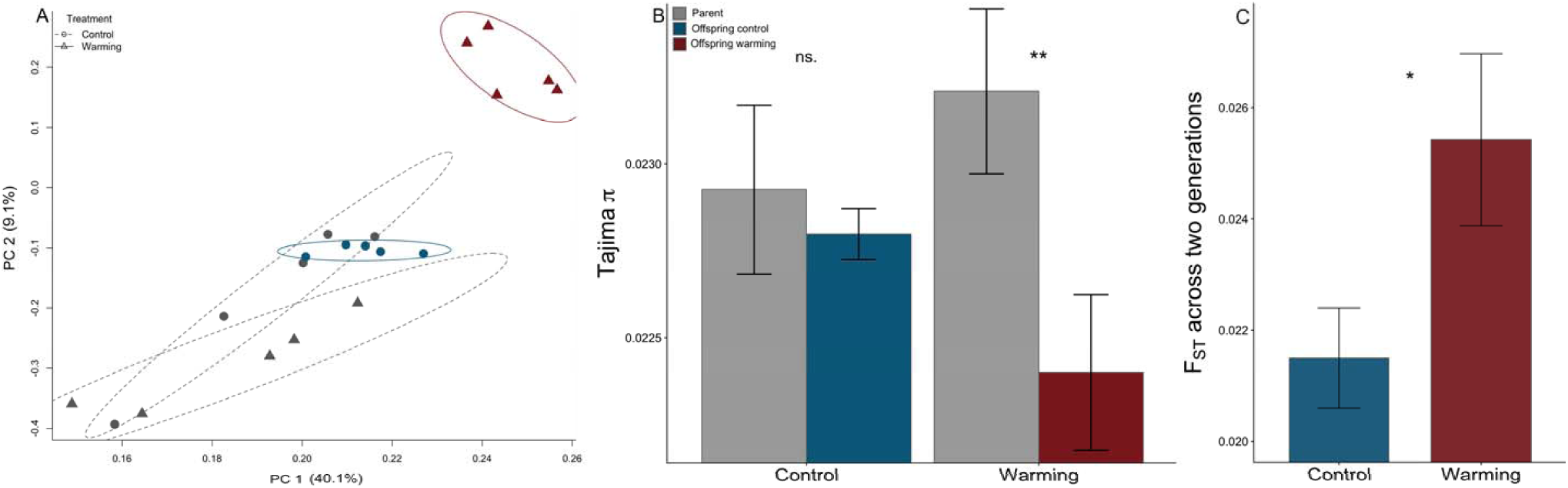
(A) Principal component analysis (PCA) visualizing genetic differences, using all polymorphic SNP markers, among the 10 experimental populations of *Ambrosia artemisiifolia*, with grey, blue and red symbols indicating parental, control offspring and warming offspring, respectively. The ellipses represent 95% confidence interval. (B) Genetic variability (Tajima’s π) within experimental *Ambrosia artemisiifolia* populations in different generations and climate change treatments. (C) Average cross-generation population differentiation (*F*_ST_) in the two treatments. All error bars are ± 1 SE. Asterisks indicate significant differences at P < 0.05 (*) or P < 0.01 (**).

We found significant variation among maternal families for all nine phenotypic traits in the parental generation (Appendix Fig. S2; all *F* ≥ 1.71, *P*_adj_ ≤ 0.03), indicating substantial genetic variation for selection to act upon. When we tested for phenotypic changes between generations and treatments, we found significant changes in days to flowering (χ^2^ = 36.98, *P*_adj_ < 0.001), the number of developing male inflorescences (χ^2^ = 13.05, *P*_adj_ < 0.001), and the total biomass of ragweed (χ^2^ = 14.17, *P*_adj_ = 0.006)(Table S1; Fig. 3). Offspring plants from the warming treatment flowered significantly later and produced larger total biomass than control offspring, or than parental plants (ghlt tukey *P* ≤ 0.005), but there were no differences between offspring control and parental plants in these traits (ghlt tukey *P* ≥ 0.85; Fig. 3A+C). In addition, offspring from warming populations had significantly more developing male inflorescences indicating larger male reproductive output than offspring from control populations (ghlt tukey *P* = 0.001), but not than parental plants (ghlt tukey *P* = 0.24; Fig. 3B). For seed traits, i.e., seed size and seed germination rate (χ^2^ ≤ 2.93, *P*_adj_ ≥ 0.18) and all other plant phenotypic traits (χ^2^ ≤ 2.73 and *P*_adj_ ≥ 0.59) (Table S1), there were no significant differences between generations and treatments. There was a significant negative correlation between total plant biomass and soil moisture content for parental plants and control offspring (both *P*_adj_ < 0.001, *r*^*2*^ ≥ 0.18), but not for warming offspring (*P*_adj_ = 0.33, *r*^*2*^ = 0.02; Fig. 4).

**Fig. 3.**
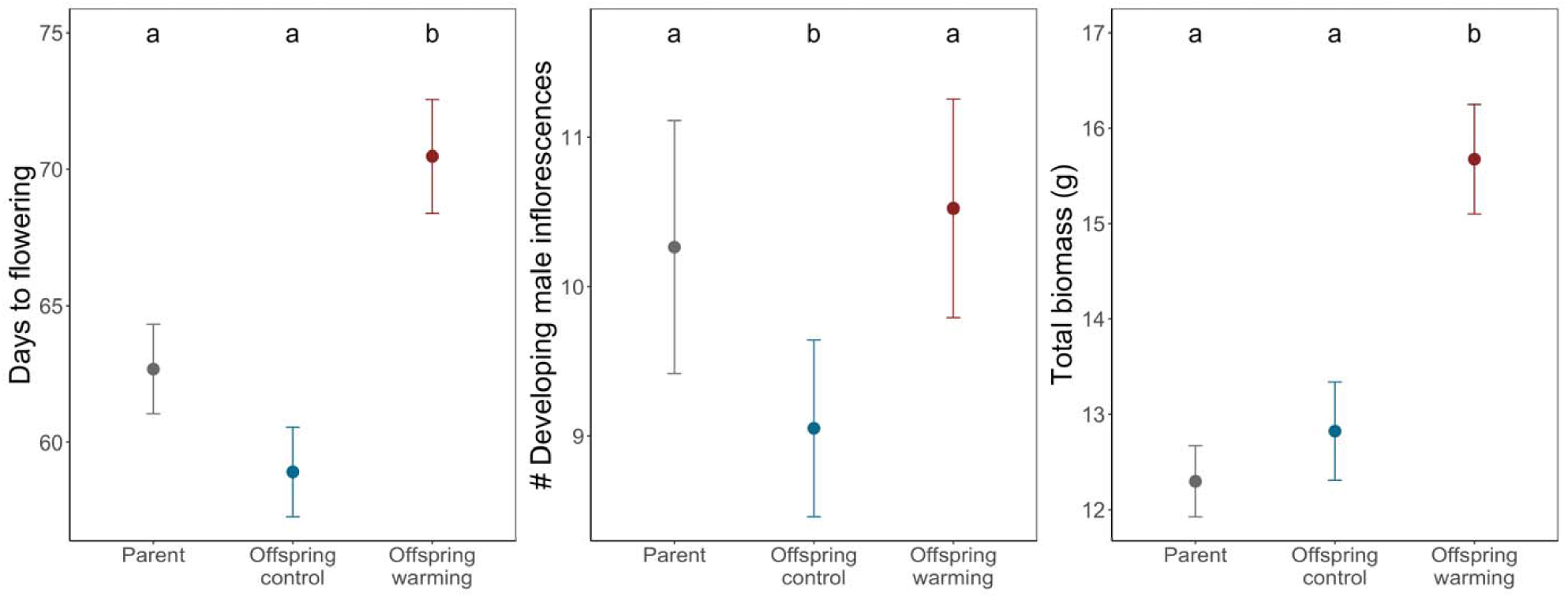
Phenotypic differences between parental and offspring plants of *Ambrosia artemisiifolia* from the two experimental treatments, when compared in a common environment. The error bars are ± 1SE. The letters on top indicate significant differences at *P*_adj_ < 0.05.

**Fig. 4.**
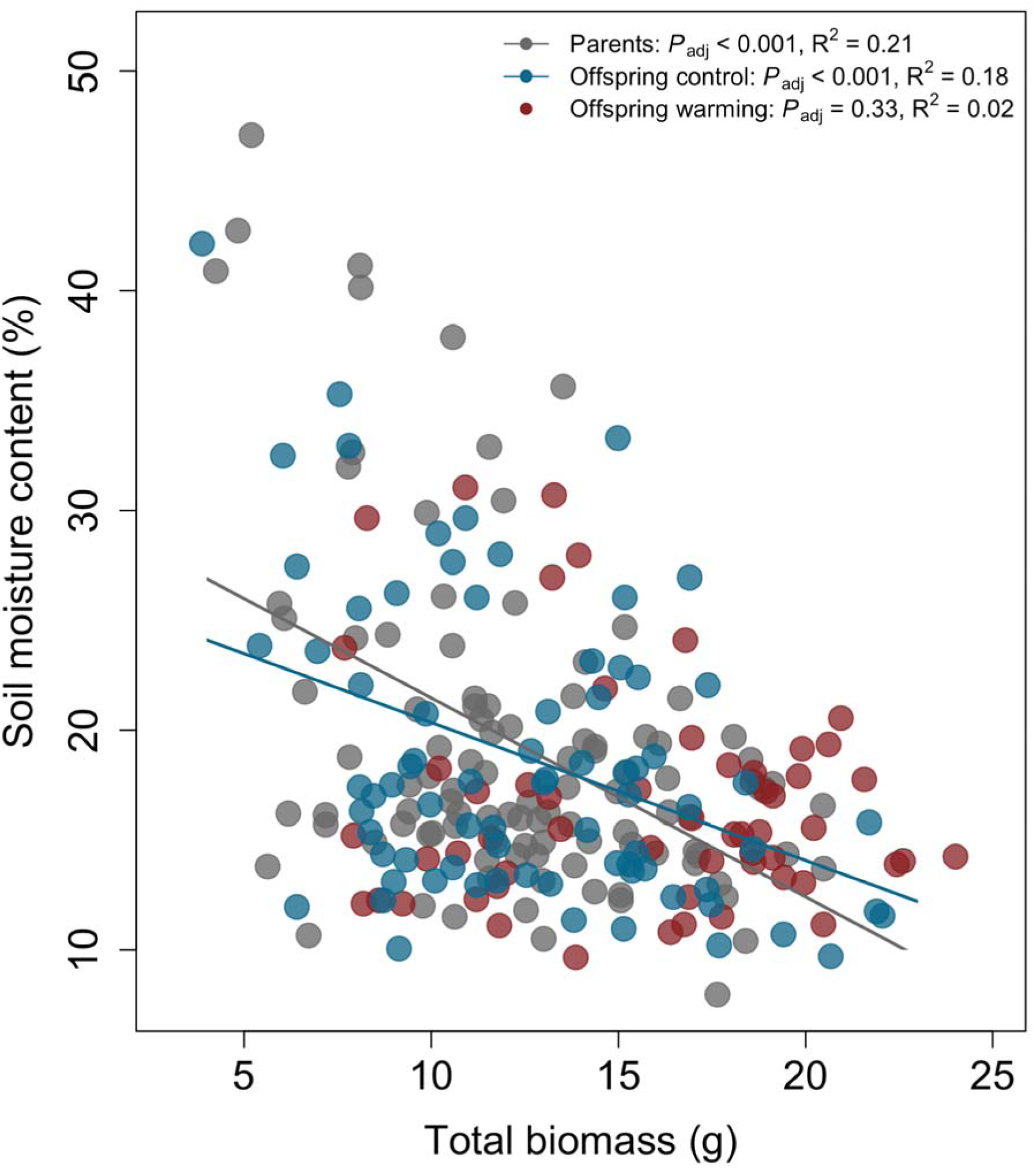
Relationships between total plant biomass and soil moisture content for parental (grey symbols) as well as for offspring plants of *Ambrosia artemisiifolia* from the control treatment (blue symbols) and from the warming treatment (red symbols). All data are from a common environment. The lines represent significant regression fits.

In seven out of nine phenotypic traits, offspring from ragweed populations that experienced warming had significantly lower *P*_ST_ value, i.e. a lower phenotypic differentiation among replicate populations, than offspring from control populations (Fig. 5A and Appendix Fig. S3). In two of the nine studied phenotypic traits – days to flowering and total biomass – we found significant differences between control and warming populations in their ratio of phenotypic to genetic differentiation, and in both cases the ratio was substantially higher for warming populations (*PG*_*W*_ > *PG*_*C*_; χ^2^ ≥ 7.82, *P*_adj_ ≤ 0.005; Fig. 5B).

**Fig. 5.**
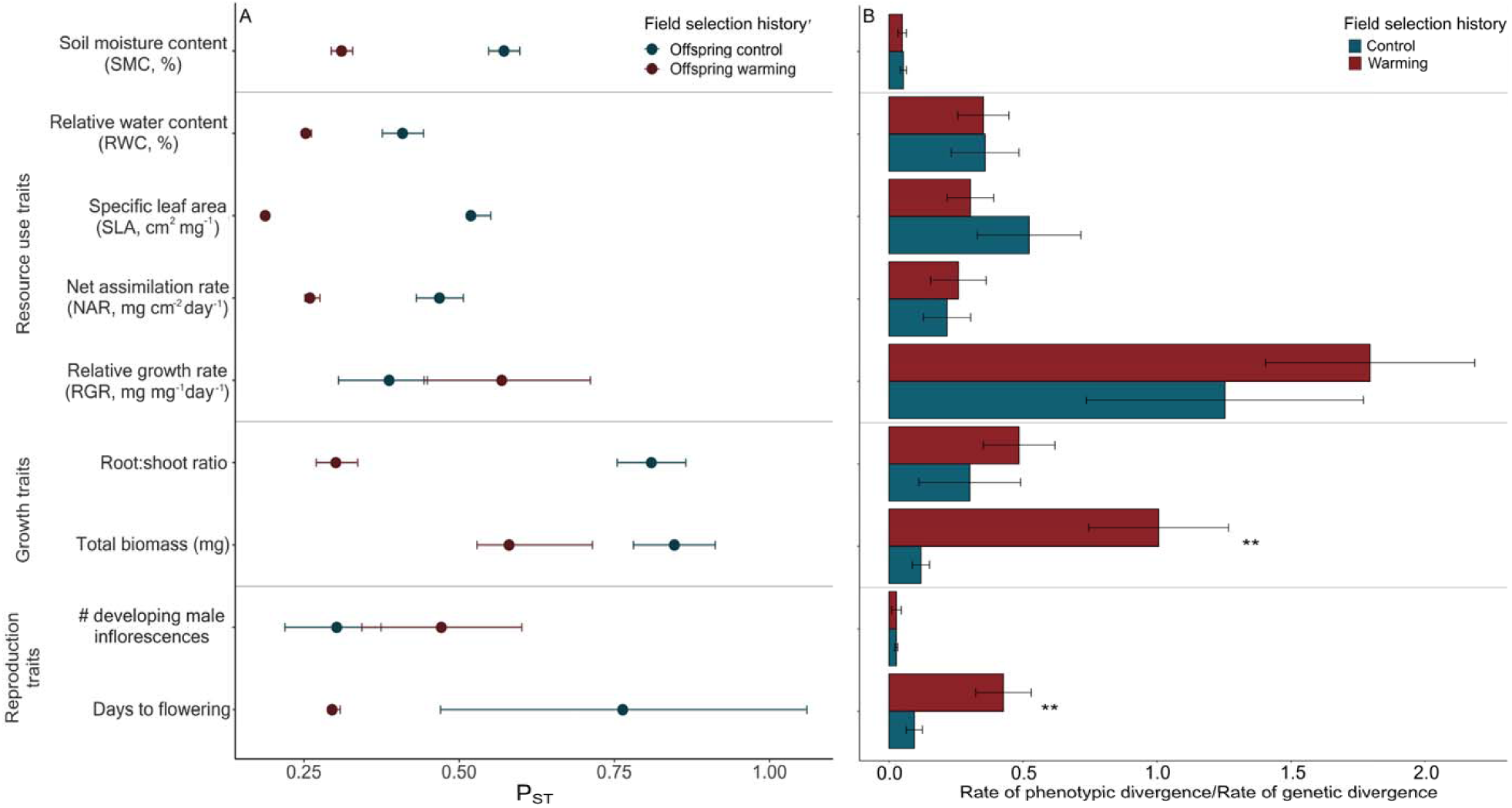
(A) Phenotypic differentiation (*P*_ST_) among experimental *Ambrosia artemisiifolia* populations from the control versus warming treatment. The values are *P*_ST_ estimates for c/*h*^2^ =1, with 99% bootstrap confidence intervals. See Fig. S3 for *P*_ST_ estimates under different c/*h*^2^ assumptions. (B) Ratios of phenotypic to genetic divergence rates across two generations for *Ambrosia artemisiifolia* populations subjected to warming versus control conditions. The error bars are ± 1SE. Asterisks indicate significant differences at *P*_adj_ < 0.01 (**).

## Discussion

Forecasting the ecological and evolutionary consequences of the interaction between plant invasion and climate change requires a thorough understanding of the underlying biological processes in plant invaders. We know from previous studies that environmental change can result in rapid selection and evolution of plant phenotypes. For instance, Skroppa and Kohmann^48^ demonstrated adaptation of Norway spruce (*Picea abies*) to local climatic conditions after only one generation, and Franks et al.^49^ found rapid evolution of flowering time in the annual *Brassica rapa* in response to climate change after just three generations. More recently, Nguyen et al.^50^ showed earlier flowering in two invasive plant species in response to selection over few generations of water manipulations, and Nowak et al.^51^ demonstrated adaptive evolution of the Alpine Pennycress *Noccaea caerulescens* after one generation of exposure to various levels of zinc contamination in the soil. Due to multiple introductions and admixtures between differentiated populations, either prior or post-introduction, invasive plants can exhibit large within-population genetic diversity^52^, This has also been found for ragweed in Europe^23,44,45^. Hahn and Rieseberg^53^ report considerable heterosis effects among particular native common ragweed population crosses, especially under simulated herbivory, but not so in crosses from the introduced French population, possibly indicating that these populations were already admixed and benefit little from further mixing. This has recently been confirmed by van Boheemen et al.^45^, who showed that historical admixture zone within native North America originated before global invasion of this weed and that European ragweed populations most likely established through multiple introductions from the native range, including those from admixed populations. This resulted in rapid adaptation of life history traits including size and flowering time to climate (latitude) in Europe^25,54^ and thus illustrates the potential for rapid adaptation to environmental changes in ragweed.

One component for such evolutionary changes in invasive species can be the increased genetic variation and of novel or transgressive phenotypes created through admixtures^11,12,25^. In addition, a small number of loci with large phenotypic effects^55^, or strong directional selection^56^ as in our study, may further contribute to rapid evolution. Here, we found significant genetic divergence in several phenotypic traits of ragweed populations under warming selection. Since both seeds collected from the parents and the second-year seedlings are purely first-generation, our results demonstrate evolutionary changes within a single generation.

The starting material for our selection experiment were 60 seed families from 19 invasive ragweed populations collected across Northern Italy. We found significant phenotypic variation among seed families for all studied traits, which corroborates previous reports of substantial genetic variation within introduced European populations^44,45^. Most importantly, it confirms that our experimental populations initially harboured plenty of raw material for selection to act upon, and that our artificial populations well reflect the high genetic within population variation described for this area^45^.

Pool-sequencing the experimental populations over two generations showed that warming populations rapidly became genetically impoverished (lower within population genetic variation in offspring as compared to parents), while this was not the case for the control populations. Moreover, in contrast to control populations, warming populations had higher *F*_ST_ values and greater genetic differentiation between parental and offspring generations. Together, these results show that our experimental warming treatment strongly accelerated evolutionary change. In addition, the populations for both the control and warming treatment became more similar within the offspring generation (smaller ellipses) as compared to the parental generation, indicating directional selection in both treatments.

The experimental populations not only evolved at the level of DNA sequence, but also at the level of phenotype. Under warming conditions, plants became larger and they flowered later, which might indicate selection towards faster growth and larger biomass accumulation, a syndrome of more efficient resource use, resulting in increased reproductive output^57^. van Boheemen et al.^25^ recently examined divergence of life-history traits in relation to climate in native North American versus introduced European and Australian populations of ragweed, and found that climate change is likely to select for larger ragweed plants that flowered later, which is in line with our results. Similarly, McGoey et al.^54^ found that larger and later-flowering ragweed plants are associated with to warmer climates and lower latitudes in both North America and Europe. It is very likely that the phenotypic changes observed in our warming treatment are also the result of natural selection. First, we found no genetic separation between generations for the control populations, and thus increased genetic drift alone cannot explain the results. Second, we found significantly higher phenotypic to genomic differentiation ratios (PG ratio) in the warming treatment than in the control treatment exactly for the two divergent traits flowering time and total biomass. However, rather than through selection for adaptive phenotypes, our findings could also be explained by the failure of a subset of maladaptive plant genotypes to produce seeds under elevated temperature conditions. The fact that these differences were apparent already after a single generation rather indicates direct effects of warming on plant physiology than indirect effects through intraspecific competition, although the warming treatment also might have increased competition for water.

Following this line of arguments, the absence of a relationships between soil moisture content (SMC) and total biomass observed only in the offspring from the warming treatment, but with a negative relationship in the parental ragweed plants and the control offspring, may then simply reflect the loss of maladapted small plants with associated high soil moisture contents (Fig. 4b). A similar effect may have caused the observed higher biomass and later flowering under warming conditions, as compared to the parental and control offspring plants, with smaller and later flowering individuals being sorted out under warming conditions.

We acknowledge that environmental maternal effects might have contributed to the observed phenotypic differentiation between treatments^58^. However, maternal effects are often found to be most pronounced during early development, i.e. during dormancy, germination and seedling growth, but to decrease over time. Effects on adult plants are often still a consequence of changes in seed size^58^. In our experiment, seed sizes and germination rates were similar for offspring from control and warming populations, indicating that maternal effects may not have played a significant role. As recommended by Moloney et al.^59^, we included plant height at transplanting as a covariate in the statistical analysis to compensate for possible maternal effects visible at that stage. In a recent study on invasive European ragweed, Gallien et al.^23^ also found that seed mass, expected to capture part of maternal effects, included as a covariable of the trait-environment regressions did not affect their results when growing plants from different altitudes in a common garden. Furthermore, they found the coefficient of genetic variation for plant height not to decrease, but to increase over time, indicating that the maternal effect was probably negligible.

In summary, the observed losses of phenotypic correlations in the warming populations may primarily result from the reduced genetic diversity^60^ due to failure of maladapted genotypes to produce seeds, which is in line with our above DNA sequencing results and the reduced *P*_ST_ for seven out of nine traits among the warming populations. Our results therefore demonstrate that climate change, as simulated by our warming treatment, can have rapid effects on the genetic composition of ragweed populations. Selection for particular genotypes may allow some traits to become more prominent in the mid- to longer-term. Such evolutionary changes were recently found to be responsible, at least partially, for niche shifts in the introduced European range of ragweed^23,25^. Moreover, larger ragweed plants, as selected for in our warming treatment, have previously been found to have higher per-capita seed and pollen production^57^. Larger plants are thus expected to further increase the future spread and impact of ragweed under changing climatic conditions.

## Conclusion

Adaptive evolution over short timescales is well-documented in invasive species^22^, specifically also for common ragweed^23,25,26,54^. Our approach combining comparisons between generations (allochronic) and between treatments (synchronic) in an experimental evolutionary field study provided a powerful test for rapid genetic responses to selection. We show that invasive ragweed populations may rapidly evolve towards larger biomass accumulation under conditions of climate warming, and that such changes can take place within a single generation. Our results also indicate that this was mainly a consequence of a rapid loss of genetic diversity, most likely through selection acting against maladapted genotypes with unfavourable physiological characteristics. Short-term evolutionary responses to climate change may aggravate the impact of some plant invaders in the future and should be considered when making predictions about future distributions and impact of plant invaders.

## Materials and Methods

Common ragweed, native to North America, has become a problematic alien invasive plant in many continents, e.g., Asia, Oceania and Europe^61^. It causes great damage to societies because of its highly allergenic pollen, and because it is also an important and hard-to-control crop weed^62^. Because of these problems, and since it is predicted to further expand in Europe under climate change, ragweed has greatly contributed to the awareness of the invasive species problem in Europe^62^.

### Field experiment

In 2016, we set up five pairs of cages (2 × 2 × 2 m) with genetically similar experimental ragweed populations in a field in Magnago, Northern Italy (Fig. 1). The municipality of Magnago, located in one of the most infested ragweed areas of Europe, provided an ideal environment for our field selection experiments in an enclosed grassland property where ragweed had been observed only very rarely before, and therefore no ragweed was expected in the soil seed bank. Each caged population was founded by 120 individuals, with two individuals from each of the same 60 maternal families that had previously been sampled from 19 invasive ragweed populations in 2013-2015 (2-4 maternal families per population; Fig. 1) in a radius of 35 km to capture the earlier described high genetic within population variation for Northern Italy^45^. Half of the experimental populations were subjected to simulated climate warming created by an open-top Plexiglas chamber, which increased the temperature, but minimized other ecological effects^63^. All plots were covered with fine-meshed tissue to protect the plants from herbivory. To verify our climate treatments, we installed temperature loggers in each experimental population and compared their measurements to the climate data in the predicted current and future ragweed distribution areas based on Sun et al.^39,40^ and climate data obtained from www.worldclim.org. The warming plots in our experiment showed a higher diurnal temperature range (χ^2^ = 4.44, *P* = 0.03), a 2.2°C increase of daily mean temperature (χ^2^ = 7.43, *P* = 0.006) and 3°C increase of daily maximum temperature (χ^2^ = 4.92, *P* = 0.02) compared to the control plots during the growing season (Appendix Fig. S1A-C). These changes were qualitatively similar to future climate changes predicted for the area suitable for ragweed during the growing season in Europe, with larger temperature ranges (*t* = -169.24, *P* < 0.001), a 3.8°C increase in monthly mean temperature (*t* = -277.02, *P* < 0.001) and a 4.8°C increase in monthly maximum temperature (*t* = -307, *P* < 0.001) (Appendix Fig. S1*D-F*). Our experimental conditions thus mimicked the predicted future climate changes reasonably well.

### Genomic analyses

To assess molecular diversity and differentiation among experimental generations and treatments, we collected leaf tissue from all 120 parental individuals in each of the ten caged field plots in Italy in June 2016, and then again from 120 offspring individuals in each plot in June 2017 (one random individual per cell in a 100 ⍰ 120 cm grid in the centre of the plot). 20 pooled populations that each contained equal amounts of each individual plant tissue (∼1mg) were then used for DNA sequencing on an Illumina HiSeq3000 at the Max Planck Institute for Developmental Biology in Tübingen (see Appendix S1C for details on DNA extraction, library preparation and processing of the raw data). We estimated genetic variability within populations as Tajima’s π with the Perl script ‘Variance-sliding.pl’ in ‘POPOOLATION’^64^. Next, we calculated the genetic differentiation (*F*_*ST*_) between each parental and offspring generation with ‘fst-sliding.pl’ in ‘POPOOLATION2’^65^, using a sliding-window approach with a window size of 100 bp and 100 bp steps to identify SNPs and genomic regions with elevated differentiation between generations. We generated position-specific nucleotide metrics with ‘bam-readcount’ (version 0.8.0, https://github.com/genome/bam-readcount), and we analysed the bam-readcounts with a custom-written Perl script to detect polymorphic sites (c. 5million SNPs were identified in each pooled population). The outputs were then used for principal component analysis (PCA) to visualize the genetic differences between the two generations for the control versus warming treatments. The results for the differentiated SNPs and their annotations will be reported elsewhere. The variation in Tajima’s π and *F*_ST_ among populations were analysed with linear mixed-effects models, using the *lmer* function in R, that included selection treatment and generation as fixed factors, and cage pairs as random effects. Pairwise comparisons were performed by least-squares means using the *emmeans* package in R.

### Phenotype assays

To assess phenotypic differences among the different experimental populations, we collected five mature seeds of each individual branch in the centre 100 ⍰ 120 cm of each parental population, and used these, together with the original seeds used to establish the parental populations, to set up a common-environment study at the University of Tübingen (see Appendix S1D for further details on growing conditions). Eventually, we were able to grow 141 plants from the parental generation (47 out of the 60 maternal families from 16 populations due to low seeds/seedlings numbers of some families, with three replicates each) and 282 offspring plants (25-30 individuals per population) in two identical growth chambers. Each plant received 60 ml tap water per day, and to avoid block effects the plants were rotated among chambers every fortnight.

Prior to the experiment, we used 70 offspring seeds to calculate the relationship between seed area and seed biomass, and we later used this formula to estimate the average seed biomass of each offspring population based on the total area of 500 seeds obtained with ImageJ (version 1.51k; http://imagej.nih.gov/ij/). We recorded germination daily over three weeks, and considered seeds as germinated when the embryo was completely uncoiled. Three days after transplanting, we measured the initial height of each plant, and we also measured the height and biomass of 30 extra seedlings and calculated their correlation to be able to non-destructively estimate the initial biomass of all plants. Because of the highly allergenic pollen of ragweed, and regional health regulations, we could not have open flowers in the growth chamber at the University of Tübingen, and this precluded growing plants first for another generation under controlled conditions to reduce maternal effects. We harvested each plant when it began to flower, i.e. the male inflorescence had reached 1 cm, which was 32-90 days after transplanting. By the end of the experiment at day 90, only nine individuals had not flowered yet and were assigned as not applicable (NA) for flowering-time analyses. As ragweed plants tended to reach their maximum height and biomass at the time of flowering^66,67^, we instead used biomass at the time of harvesting as a fitness proxy, as biomass was found to be highly correlated with per-capita seed and pollen production in a field study across 39 sites in Europe^57^. At the final harvest, we counted the numbers of developing male inflorescences of each plant as a proxy for potential male reproductive output, and we measured soil moisture content (SMC) using a soil moisture sensor (Delta-T Devices, Cambridge, UK) in all pots to assess differences in water use of the plants. We then separated all plants into aboveground and root biomass, dried them at 60°C for 72 h and weighed them. To further characterise the phenotypic differences in functional traits among the plants, we also calculated NAR, the relative growth rate (RGR), RWC and SLA of each plant (see Appendix S1D for details).

We analysed our data with (generalized) linear mixed models, using the *glmer*/*lmer* functions in the R package *lme4*, which uses maximum likelihood to estimate model parameters^68^. Days to flowering and number developing male inflorescences were analysed with *glmer* using a log-link and a Poisson error distribution, and all other (normally distributed) variables – seed size, germination rate, plant biomass, RWC, SLA, RGR, NAR and SMC – were analysed with *lmer*. The models included generation and selection treatments (i.e., parent, control offspring and warming offspring) as a fixed factor, initial height as a covariate to partially account for maternal effects, and cage pairs (in the field experiment) as random effect. We adjusted *P*‐values using the Bonferroni‐Holm method to correct for type 1 error. In cases where significant differences occurred (*P*_adj_ < 0.05), we carried out Tukey’s post-hoc tests using the *glht* function in the R package *multcomp*^69^ to compare parents and the two offspring treatments. To further analyse correlations between different traits, we used mixed-effects regression models in which cage pairs were again treated as random effects.

To quantify genetically-based phenotypic differentiation among populations (P_*ST*_) we used the *Pst* function in the R package *Pstat*^70^, with 99% confidence intervals from 5000 bootstrapped estimates. How well *P*_ST_ approximates Q_*ST*_ depends on the heritability *h*^2^ and the between-population additive genetic component *c*^71^. Following Sun and Roderick^72^, we set heritability to 0.46-0.95 and c from 0.01 (1% phenotypic variance due to additive genetic effects) to 1 (all phenotypic variance due to additive genetic effects) and calculated *P*_ST_ for this range of *h*^*2*^ and *c* values. To test a realistic range of possibilities, we thus calculated *P*_ST_ for five values of c/*h*^2^ (0.01, 0.5, 1, 1.5, 2).

### Comparisons of phenotypic and genetic divergence

In microevolutionary studies, phenotypic and genetic analyses provide different, but complementary information. Specifying a change as phenotypic does not imply that the change itself was not genetic, but simply that the relative contribution of genetic and non-genetic effects is not known^73^. Phenotypic and genetic changes can be assessed allochronically when comparing the same population at different points in time (i.e., temporal context; between generations in our study), or synchronically by comparing populations that had a common origin in the past (i.e., spatial context or space-for-time approach; between treatments within generations in our study)^74,75^. Allochronic studies are used to estimate the pace of evolutionary change, whereas synchronic studies more appropriately estimate divergence^73^. In the synchronic approach, different populations can be grown together under specific conditions, while an allochronic approach is generally harder to implement, or sometimes impossible^74^. A combined approach will allow assessing the population differentiation in time and space simultaneously^76^. To infer the adaptive evolution of phenotypic changes in response to climate change, we can think of a synchronic comparison to assess if the rate or directionality of evolutionary change in the selection treatment (allochronic setting) exceeds the one in the control treatment (cf. Fig. 1). This approach has been applied in our study.

To estimate the effects of our experimental treatment on molecular and phenotypic changes across generations (i.e. rates of divergence), and to test whether these data are consistent with a hypothesis of rapid response to selection, we first estimated, in the allochronic comparisons, the rates of phenotypic and sequence divergence, separately for each experimental population and trait. Including the allochronic comparison in our analysis considers the fact that the initial populations were similar (same maternal families), but not identical due to high to complete outcrossing of ragweed^27,43^. In a second step, we then calculated the ratios between the rates of phenotypic and sequence divergence. More details on how we estimated phenotypic divergence (*P*_*C*_ and *P*_*W*_ for control and warming; respectively, Fig. 1) and sequence divergence (*G*_*C*_ and *G*_*W*_ for control and warming; respectively, Fig. 1) and the ratio between them (*PG*_*C*_ and *PG*_*W*_ for control and warming; respectively) are given in Appendix S1E. A larger value of the ratio will indicate stronger selection on phenotypes^74^, and the comparison between the ratio with and without warming selection can thus infer us about the relative strength of natural selection in the experimental warming treatments. We expected that *PG*_*W*_ would be larger than *PG*_*C*_ if the simulated warming caused indeed stronger genetic response to selection than under control. We calculated and analysed the variation in the ratios with linear mixed models that included the selection treatment as a fixed factor, and cage pairs as random effects; *P*‐values were then adjusted to correct for type 1 errors. All statistical analyses were done in R version 3.4.3 (R Development Core Team, 2017), and all figures created with the R package *ggplot2*.

## Acknowledgements

YS was supported by an Advanced Postdoc.Mobility fellowship from the Swiss National Science Foundation (SNSF; Project No. P300PA_161014), with additional support from the Novartis Foundation (#17B083 to HMS and YS). YS and OB acknowledge support by the High Performance and Cloud Computing Group at the Zentrum für Datenverarbeitung of the University of Tübingen, the state of Baden-Württemberg through bwHPC and the German Research Foundation (DFG) through grant no INST 37/935-1 FUGG. HMS acknowledges funding through the Swiss National Science Foundation (project number 31003A_166448). We gratefully acknowledge the support and help of Detlef Weigel, Gautam Shirsekar, Julia Hildebrandt and Ilja Bezrukov with the DNA sequencing, Martin Kapun for their help with the pool-seq analyses and Daniel Wegmann and Jérôme Goudet for their suggestions on how to assess selection through ratios of phenotypic and genetic divergences in our study. We also thank Kay Hodgins and Michael Martin for the draft genome of *Ambrosia artemisiifolia*.

## Date availability statement

Data associate with the manuscript are deposited in Fig-share Repository: http://doi.org/10.6084/m9.figshare.9994415.

## Appendix S1: Supplementary methods

A. *Field Temperature treatments* and statistical analyses
B. *DNA extraction and genome sequencing*
C. *Illumina read processing, mapping and SNP calling*
D. *Growth chamber experiment for phenotypic assays*
E. *Ratios of phenotypic and genetic divergence*

### Appendix: supplementary table and figures

Table S1. Statistical comparisons of phenotypic trait means between parental and offspring plants from the two experimental treatments.

Fig. S1. Average differences in climatic conditions between the treatments in our field experiment (A-C) and current vs. predicted future climates for the distribution range of *Ambrosia artemisiifolia* from the species distribution models (D-F). (A) Diurnal temperature ranges, (B) daily mean temperatures, and (C) daily maximum temperatures in the control (blue) and warming (red) treatment during the growing season from April to October. (D) Monthly temperature differences (April – October), (E) mean temperatures of the warmest quarter of the year, and (F) monthly maximum temperatures for the current climate (blue) and predicted future climate in the distribution range of *A. artemisiifolia* (see Appendix S1A for details on data sources). The vertical dashed lines mark the respective mean values.

Fig. S2. Phenotypic traits of the *Ambrosia artemisiifolia* parental generation plants assessed under control conditions.

Fig. S3. Estimated phenotypic differentiation (*P*_ST_) among experimental *Ambrosia artemisiifolia* populations under control versus warming conditions, tested for a range of *c/h*^*2*^ ratios (*c* = strength of additive genetic effects, *h*^*2*^ = heritability) in nine quantitative traits. Error bars are 99% bootstrap confidence limits.

## References

1. IPCC. Climate change 2013: the physical science basis. Contribution of working group I to the fifth assessment report of the intergovernmental panel on climate change. (eds Stocker TF, et al.). Cambridge University Press (2013).

2. Chevin L-M, Hoffmann AA. Evolution of phenotypic plasticity in extreme environments. Phil Trans R Soc B 372, 20160138 (2017).

3. Alberto FJ, et al. Potential for evolutionary responses to climate change–evidence from tree populations. Global Change Biol 19, 1645–1661 (2013).

4. Lavergne S, Mouquet N, Thuiller W, Ronce O. Biodiversity and climate change: integrating evolutionary and ecological responses of species and communities. Annu Rev Ecol, Evol Syst 41, 321–350 (2010).

5. Merow C, Bois ST, Allen JM, Xie Y, Silander JA. Climate change both facilitates and inhibits invasive plant ranges in New England. Proc Natl Acad Sci 114, E3276–E3284 (2017).

6. Seebens H, et al. No saturation in the accumulation of alien species worldwide. Nat Commun 8, 14435 (2017).

7. Vilà M, et al. Ecological impacts of invasive alien plants: a meta-analysis of their effects on species, communities and ecosystems. Ecol Lett 14, 702–708 (2011).

8. Sandel B, Dangremond EM. Climate change and the invasion of California by grasses. Global Change Biol 18, 277–289 (2012).

9. Alexander JM, Edwards PJ. Limits to the niche and range margins of alien species. Oikos 119, 1377–1386 (2010).

10. Dlugosch KM, Anderson SR, Braasch J, Cang FA, Gillette HD. The devil is in the details: genetic variation in introduced populations and its contributions to invasion. Mol Ecol 24, 2095–2111 (2015).

11. Rius M, Darling JA. How important is intraspecific genetic admixture to the success of colonising populations? Trends Ecol Evol 29, 233–242 (2014).

12. Bock DG, et al. What we still don’t know about invasion genetics. Mol Ecol 24, 2277–2297 (2015).

13. Strauss SY. Ecological and evolutionary responses in complex communities: implications for invasions and eco-evolutionary feedbacks. Oikos 123, 257–266 (2014).

14. Bone E, Farres A. Trends and rates of microevolution in plants. Genetica 112, 165–182 (2001).

15. Chown SL, Hodgins KA, Griffin PC, Oakeshott JG, Byrne M, Hoffmann AA. Biological invasions, climate change and genomics. Evol Appl 8, 23–46 (2015).

16. Ravenscroft CH, Whitlock R, Fridley JD. Rapid genetic divergence in response to 15 years of simulated climate change. Global Change Biol 21, 4165–4176 (2015).

17. Hoffmann AA, Sgrò CM. Climate change and evolutionary adaptation. Nature 470, 479–485 (2011).

18. Olsson K, Ågren J. Latitudinal population differentiation in phenology, life history and flower morphology in the perennial herb Lythrum salicaria. J Evol Biol 15, 983–996 (2002).

19. Colautti RI, Ågren J, Anderson JT. Phenological shifts of native and invasive species under climate change: insights from the Boechera–Lythrum model. Philos Trans R Soc B: Biol Sci 372, 20160032 (2017).

20. Li J, Du L, Guan W, Yu F-H, van Kleunen M. Latitudinal and longitudinal clines of phenotypic plasticity in the invasive herb Solidago canadensis in China. Oecologia 182, 755–764 (2016).

21. Moran EV, Alexander JM. Evolutionary responses to global change: lessons from invasive species. Ecol Lett 17, 637–649 (2014).

22. Prentis PJ, Wilson JRU, Dormontt EE, Richardson DM, Lowe AJ. Adaptive evolution in invasive species. Trends Plant Sci 13, 288–294 (2008).

23. Gallien L, et al. Is there any evidence for rapid, genetically-based, climatic niche expansion in the invasive common ragweed? PloS one 11, e0152867 (2016).

24. Coltman DW, O’donoghue P, Jorgenson JT, Hogg JT, Strobeck C, Festa-Bianchet M. Undesirable evolutionary consequences of trophy hunting. Nature 426, 655–658 (2003).

25. van Boheemen LA, Atwater DZ, Hodgins KA. Rapid and repeated local adaptation to climate in an invasive plant. New Phytol, in press (2018).

26. Chun YJ, Le Corre V, Bretagnolle F. Adaptive divergence for a fitness-related trait among invasive Ambrosia artemisiifolia populations in France. Mol Ecol 20, 1378–1388 (2011).

27. Li X-M, Liao W-J, Wolfe LM, Zhang D-Y. No evolutionary shift in the mating system of North American Ambrosia artemisiifolia (Asteraceae) following its introduction to China. PloS one 7, e31935 (2012).

28. Agrawal AA, Hastings AP, Johnson MTJ, Maron JL, Salminen JP. Insect herbivores drive real-time ecological and evolutionary change in plant populations. Science 338, 113–116 (2012).

29. Schlötterer C, Kofler R, Versace E, Tobler R, Franssen S. Combining experimental evolution with next-generation sequencing: a powerful tool to study adaptation from standing genetic variation. Heredity 114, 431–440 (2015).

30. de Villemereuil P, Gaggiotti OE, Mouterde M, Till-Bottraud I. Common garden experiments in the genomic era: new perspectives and opportunities. Heredity 116, 249–254 (2016).

31. Barghi N, et al. Genetic redundancy fuels polygenic adaptation in Drosophila. PLoS Biol 17, e3000128 (2019).

32. Müller-Schärer H, Bouchemousse S, Litto M, McEvoy PB, Roderick GK, Sun Y. How to better predict long-term benefits and risks in weed biocontrol: an evolutionary perspective. Curr Opin Insect Sci, (in press).

33. Mousseau M. At the crossroads of plant physiology and ecology. Trends Plant Sci 4, 1 (1999).

34. Hudson J, Henry G, Cornwell W. Taller and larger: shifts in Arctic tundra leaf traits after 16 years of experimental warming. Global Change Biol 17, 1013–1021 (2011).

35. Barrs H, Weatherley P. A re-examination of the relative turgidity technique for estimating water deficits in leaves. Aust J Biol Sci 15, 413–428 (1962).

36. Maisto G, Santorufo L, Arena C. Heavy metal accumulation in leaves affects physiological performance and litter quality of Quercus ilex L. J Plant Nutr Soil Sci 176, 776–784 (2013).

37. Wahid A. Physiological implications of metabolite biosynthesis for net assimilation and heat-stress tolerance of sugarcane (Saccharum officinarum) sprouts. J Plant Res 120, 219–228 (2007).

38. Schaffner U, et al. Biological weed control to relieve millions of allergy sufferers in Europe. Nat Commun accepted, (2020).

39. Sun Y, Brönnimann O, Roderick GK, Poltavsky A, Lommen ST, Müller-Schärer H. Climatic suitability ranking of biological control candidates: a biogeographic approach for ragweed management in Europe. Ecosphere 8, e01731. 01710.01002/ecs01732.01731 (2017).

40. Sun Y, Zhou Z, Wang R, Müller-Schärer H. Biological control opportunities of ragweed are predicted to decrease with climate change in East Asia. Biodivers Sci 25, 1285–1284 (2018).

41. Case MJ, Stinson KA. Climate change impacts on the distribution of the allergenic plant, common ragweed (Ambrosia artemisiifolia) in the eastern United States. PloS one 13, e0205677 (2018).

42. Polechová J, Barton N, Marion G. Species’ range: adaptation in space and time. Am Nat 174, E186–E204 (2009).

43. Friedman J, Barrett SC. High outcrossing in the annual colonizing species Ambrosia artemisiifolia (Asteraceae). Annals of Botany 101, 1303–1309 (2008).

44. Genton BJ, Shykoff JA, Giraud T. High genetic diversity in French invasive populations of common ragweed, Ambrosia artemisiifolia, as a result of multiple sources of introduction. Mol Ecol 14, 4275–4285 (2005).

45. van Boheemen LA, Lombaert E, Nurkowski KA, Gauffre B, Rieseberg LH, Hodgins KA. Multiple introductions, admixture and bridgehead invasion characterize the introduction history of Ambrosia artemisiifolia in Europe and Australia. Mol Ecol 26, 5421–5434 (2017).

46. Gaudeul M, Giraud T, Kiss L, Shykoff JA. Nuclear and chloroplast microsatellites show multiple introductions in the worldwide invasion history of common ragweed, Ambrosia artemisiifolia. PLoS ONE 6, e17658 (2011).

47. Colautti RI, Lau JA. Contemporary evolution during invasion: evidence for differentiation, natural selection, and local adaptation. Mol Ecol 24, 1999–2017 (2015).

48. Skroppa T, Kohmann K. Adaptation to local conditions after one generation in Norway spruce. Int J For Genet 4, 171–177 (1997).

49. Franks SJ, Sim S, Weis AE. Rapid evolution of flowering time by an annual plant in response to a climate fluctuation. Proc Natl Acad Sci 104, 1278–1282 (2007).

50. Nguyen MA, Ortega AE, Nguyen KQ, Kimball S, Goulden ML, Funk JL. Evolutionary responses of invasive grass species to variation in precipitation and soil nitrogen. J Ecol 104, 979–986 (2016).

51. Nowak J, et al. Can zinc pollution of soil promote adaptive evolution in plants? Insights from a one-generation selection experiment. J Exp Bot 69, 5561–5572 (2018).

52. Lavergne S, Molofsky J. Increased genetic variation and evolutionary potential drive the success of an invasive grass. Proc Natl Acad Sci 104, 3883–3888 (2007).

53. Hahn MA, Rieseberg LH. Genetic admixture and heterosis may enhance the invasiveness of common ragweed. Evol Appl 10, 241–250 (2017).

54. McGoey BV, Hodgins KA, Stinchcombe JR. Parallel clines in native and introduced ragweed populations are likely due to adaptation. bioRxiv, 773838 (2019).

55. Lendvai G, Levin D. Rapid response to artificial selection on flower size in Phlox. Heredity 90, 336–342 (2003).

56. Rego A, Messina FJ, Gompert Z. Dynamics of genomic change during evolutionary rescue in the seed beetle Callosobruchus maculatus. Mol Ecol 28, 2136–2154 (2019).

57. Lommen STE, et al. Explaining variability in the production of seed and allergenic pollen by invasive Ambrosia artemisiifolia across Europe. Biol Invasions 20, 1475–1491 (2017).

58. Roach DA, Wulff RD. Maternal effects in plants. Annu Rev Ecol Syst 18, 209–235 (1987).

59. Moloney KA, Holzapfel C, Tielbörger K, Jeltsch F, Schurr FM. Rethinking the common garden in invasion research. Perspect Plant Ecol Evol Syst 11, 311–320 (2009).

60. Roff D. The evolution of the G matrix: selection or drift? Heredity 84, 135–142 (2000).

61. Essl F, et al. Biological Flora of the British Isles: Ambrosia artemisiifolia. J Ecol 103, 1069–1098 (2015).

62. Müller-Schärer H, et al. Cross-fertilizing weed science and plant invasion science to improve efficient management: A European challenge. Basic Appl Ecol 33, 1–13 (2018).

63. Marion G, et al. Open-top designs for manipulating field temperature in high-latitude ecosystems. Global Change Biol 3, 20–32 (1997).

64. Kofler R, et al. PoPoolation: a toolbox for population genetic analysis of next generation sequencing data from pooled individuals. PloS one 6, e15925 (2011).

65. Kofler R, Pandey RV, Schlötterer C. PoPoolation2: identifying differentiation between populations using sequencing of pooled DNA samples (Pool-Seq). Bioinformatics 27, 3435–3436 (2011).

66. Sun S, Frelich LE. Flowering phenology and height growth pattern are associated with maximum plant height, relative growth rate and stem tissue mass density in herbaceous grassland species. J Ecol 99, 991–1000 (2011).

67. Lommen ST, et al. Time to cut: population models reveal how to mow invasive common ragweed cost-effectively. NeoBiota 39, 53–78 (2018).

68. Bates D, et al. Package ‘lme4’. R foundation for statistical computing, Vienna 12, (2014).

69. Hothorn T, Bretz F, Westfall P. Simultaneous inference in general parametric models. Biom J 50, 346–363 (2008).

70. Da Silva SB, Da Silva A. Pstat: an R Package to Assess Population Differentiation in Phenotypic Traits. R J 10, 447–454 (2018).

71. Brommer J. Whither PST? The approximation of QST by PST in evolutionary and conservation biology. J Evol Biol 24, 1160–1168 (2011).

72. Sun Y, Roderick GK. Rapid evolution of invasive traits facilitates the invasion of common ragweed, Ambrosia artemisiifolia. J Ecol 107, 2673–2687 (2019).

73. Hendry AP, Kinnison MT. Perspective: the pace of modern life: measuring rates of contemporary microevolution. Evolution 53, 1637–1653 (1999).

74. Merilä J, Hendry AP. Climate change, adaptation, and phenotypic plasticity: the problem and the evidence. Evol Appl 7, 1–14 (2014).

75. Verheyen J, Tüzün N, Stoks R. Using natural laboratories to study evolution to global warming: contrasting altitudinal, latitudinal, and urbanization gradients. Curr Opin Insect Sci 35, 10–19 (2019).

76. Van Dijk H, Hautekèete NC. Evidence of genetic change in the flowering phenology of sea beets along a latitudinal cline within two decades. J Evol Biol 27, 1572–1581 (2014).

