## appendix for "High standing genetic variation in an invasive plant allows immediate evolutionary response to climate warming"

**Appendix: Supplementary table and figures.**

**Table S1**. Statistical comparisons of phenotypic trait of *Ambrosia artemisiifolia* among parent, control offspring and warming offspring.

| Traits | Parents  (Mean ± SE) | Offspring Control  (Mean ± SE) | Offspring Warming  (Mean ± SE) | Chi-square | *Adjusted  P*-value |
| --- | --- | --- | --- | --- | --- |
| Seed size [mg] | 5.14 ± 0.19 | 5.64 ± 0.1 | 5.35 ± 0.11 | 2.93 | 0.18 |
| Seed germination rate | 0.39 ± 0.03 | 0.45 ± 0.02 | 0.41 ± 0.06 | 0.85 | 0.36 |
| Days to flowering [days] | 62.88 ± 1.59 | 59.05 ± 1.62 | 70.48 ± 2.08 | 36.98 | **< 0.001** |
| Number of flowering buds | 10.26 ± 0.85 | 9.05 ± 5.20 | 10.53 ± 0.73 | 13.05 | **< 0.001** |
| Total biomass [g] | 12.41 ± 0.37 | 12.81 ± 0.51 | 15.68 ± 0.57 | 14.17 | **0.006** |
| Root:shoot ratio | 0.37 ± 0.02 | 0.38 ± 0.03 | 0.37 ± 0.03 | 0.19 | 0.80 |
| Relative growth rate | 0.10 ± 0.00 | 0.11 ± 0.00 | 0.09 ± 0.00 | 0.24 | 0.79 |
| Relative water content | 0.74 ± 0.01 | 0.74 ± 0.01 | 0.73 ± 0.01 | 0.68 | 0.80 |
| Specific leaf area [cm^2^ · mg^-1^] | 0.29 ± 0.01 | 0.30 ± 0.01 | 0.28 ± 0.01 | 2.73 | 0.59 |
| Net assimilation rate [mg · cm^-2^ · day^-1^] | 0.06 ± 0.00 | 0.06 ± 0.00 | 0.06 ± 0.00 | 1.18 | 0.72 |
| Soil moisture content [%] | 19.29 ± 0.73 | 18.59 ± 0.77 | 18.37 ± 1.05 | 0.01 | 0.94 |

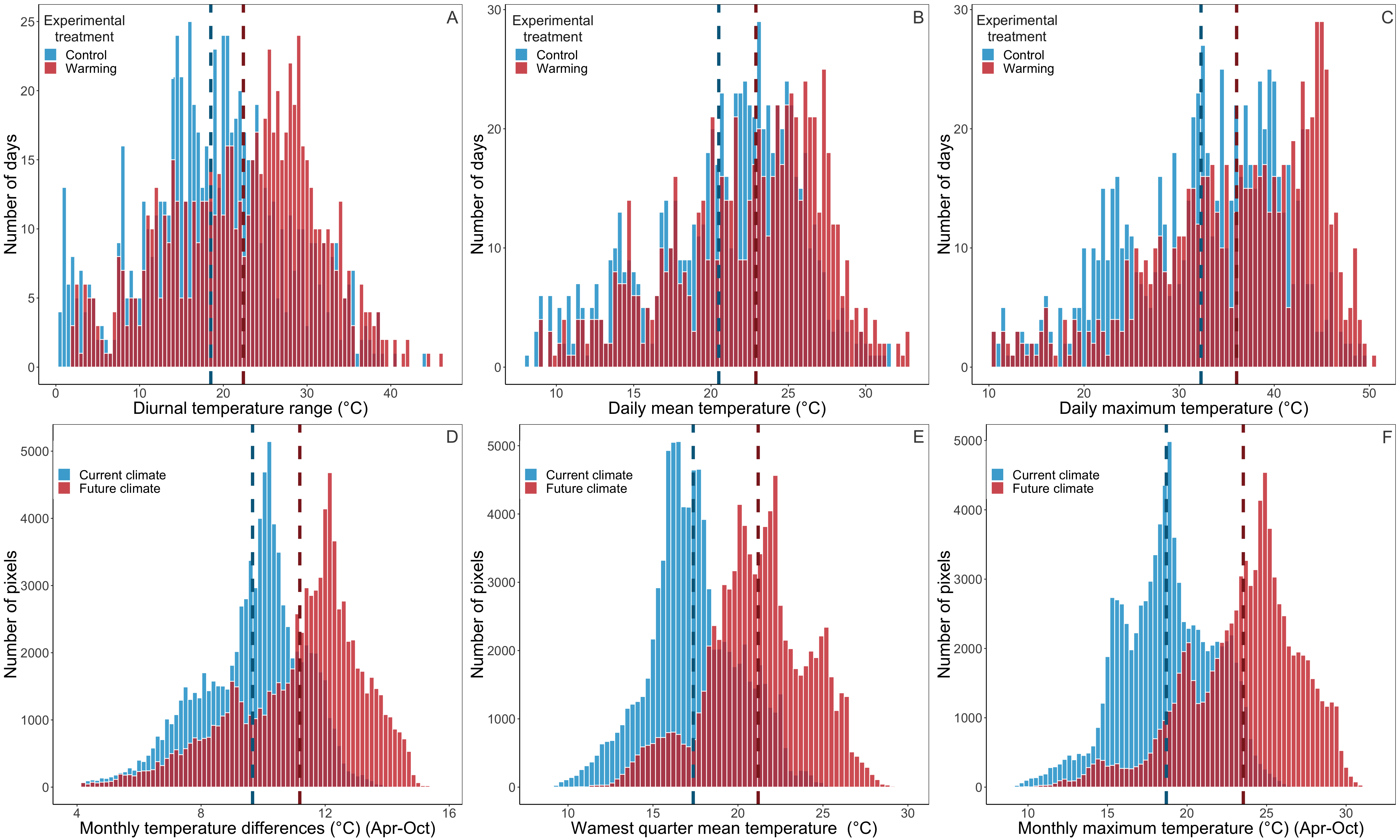

**Fig. S1.** Average differences in climatic conditions between the treatments in our field experiment (A-C) and current vs. predicted future climates for the distribution range of *Ambrosia artemisiifolia* from the species distribution models (D-F). (A) Diurnal temperature ranges, (B) daily mean temperatures, and (C) daily maximum temperatures in the control (blue) and warming (red) treatment during the growing season from April to October. (D) Monthly temperature differences (April – October), (E) mean temperatures of the warmest quarter of the year, and (F) monthly maximum temperatures for the current climate (blue) and predicted future climate in the distribution range of *A. artemisiifolia* (see Appendix S1A for details on data sources). The vertical dashed lines mark the respective mean values.

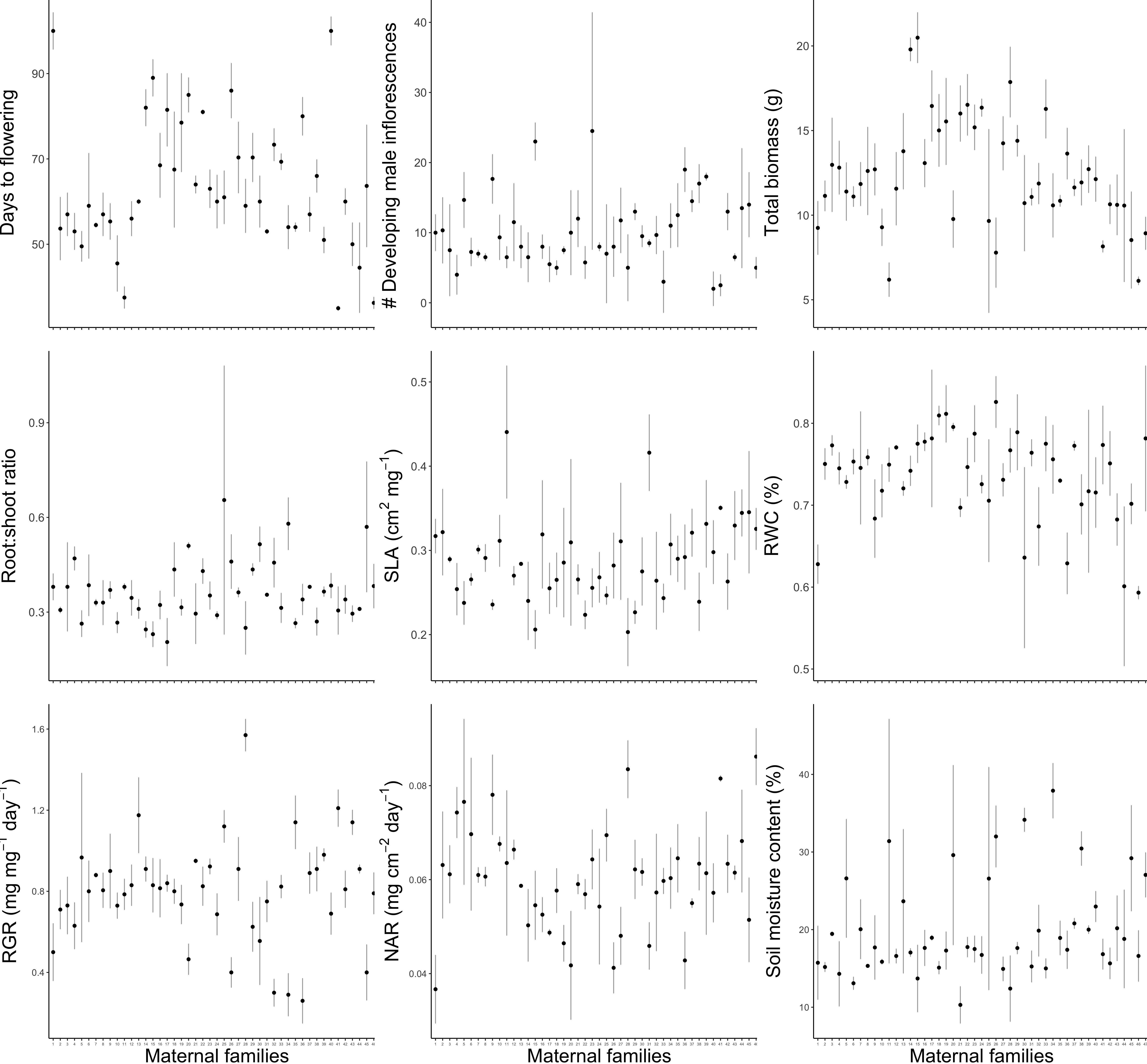

**Fig. S2.** Phenotypic variation among *Ambrosia artemisiifolia* maternal families used in our experiment parental generation plants assessed under control conditions; error bars represent ± SE.

Note: to explore the initial genetic variation across all families within parental-generation-only, we used the linear model and generalized linear models, using the lm/*glm* function, for normal and Poisson destruction data, respectively.

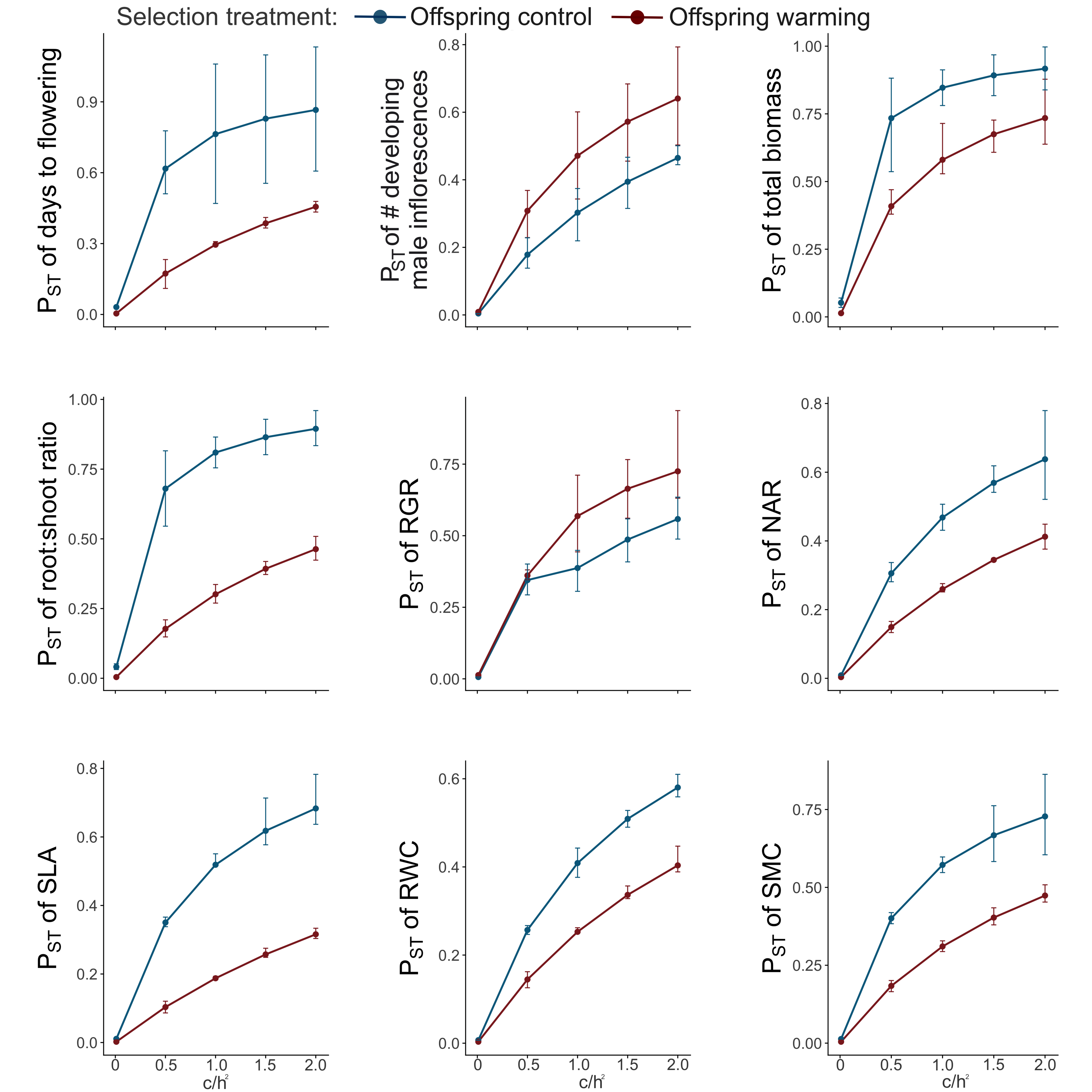
**Fig. S3.** Estimated phenotypic differentiation (*P*_ST_) among experimental *Ambrosia artemisiifolia* populations under control versus warming conditions, tested for a range of *c/h^2^* ratios (*c* = strength of additive genetic effects, *h^2^* = heritability) in nine quantitative traits. Error bars are 99% bootstrap confidence limits.
